# Polyglutamine expansion induced dynamic misfolding of Androgen Receptor

**DOI:** 10.1101/2024.12.19.629423

**Authors:** Laurens W.H.J. Heling, Vahid Sheikhhassani, Julian Ng, Morris van Vliet, Alba Jiménez-Panizo, Andrea Alegre-Martí, Jaie Woodard, Willeke van Roon-Mom, Iain J McEwan, Eva Estébanez-Perpiñá, Alireza Mashaghi

## Abstract

Spinal bulbar muscular atrophy (SBMA) is caused by a polyglutamine expansion (pQe) in the N-terminal transactivation domain of human androgen receptor (AR-NTD), resulting in a combination of toxic gain- and loss-of-function mechanisms. The structural basis of these processes has not been resolved due to the disordered nature of the NTD, which hinders experimental analyses of its detailed conformations. Here, using extensive computational modelling, we show that AR-NTD forms dynamic compact regions, which upon pQe re-organize dynamically, mediated partly by direct pQ interaction with the Androgen N-Terminal Signature (ANTS) motif. The altered dynamics of the NTD result in a perturbation of interdomain interactions, with potential implications for binding of the receptor protein to its response element. Oligomeric aggregation of the dynamic misfolded NTD exposes pQe, but blocks tau-5 and the FQNLF motif, which could lead to aberrant receptor transcriptional activity. These observations suggest a structural mechanism for AR dysfunction in SBMA.

## Introduction

The Androgen Receptor (AR/NR3C4) is a crucial ligand-activated transcription factor, which is mutated in several human pathologies^1^. Spinal Bulbar Muscular Atrophy (SBMA)^2,3^, also known as Kennedy’s Disease, is a neuromuscular condition that affects approximately 1 in 40,000 men ^4^ and is characterized by the progressive loss of motor neurons in the brainstem and spinal cord, leading to muscle atrophy and weakness in bulbar and extremity muscles^5^. The toxicity of AR in SBMA is dependent on the presence of androgens, testosterone, and its more potent derivative, dihydrotestosterone, while alterations in cellular processes ultimately lead to cell dysfunction and cell death^6–11^. The onset and progression of SBMA is associated with an expansion of the CAG trinucleotide repeat in that part of the gene that encodes the N-terminal transactivation domain (NTD), the intrinsically disordered protein (IDP) region of AR. This longer CAG repeat is translated in an expanded polyglutamine (pQ) stretch (henceforth referred to as pQe) in the protein but the molecular mechanisms and cellular pathways that lead to neuronal dysfunction and cell death remain poorly understood. For SBMA and other AR-related pathologies the underlying mechanisms remain unclear, in part, due to the limited structural models of AR.

AR is a member of the nuclear receptor superfamily^12^ and displays a distinct modular structure consisting of three structural domains (Figure 1A). While the ligand binding domain (LBD)^13–15^ and DNA binding domain (DBD)^16^ have been structurally characterized to atomic level as isolated modules, a high-resolution experimental structure of the NTD and full-length AR remains elusive, even though low-resolution models are available^17^. In a recent computational study, we presented a first insight into a high-resolution atomistic model of wild-type (wt) AR-NTD^18^. Our study revealed that the AR-NTD forms into two spatially segregated 3D regions, the N-terminal Region (NR) (residues 1-224) and C-terminal Region (CR) (225-538) (Figure 1B). The overall shape and orientation of the computational model agreed well with the cryo-EM image of the AR^17^. This work provided a first 3D conformational model of the NTD, paving the way for the analysis of native and mutated forms of AR.

**Figure 1.**
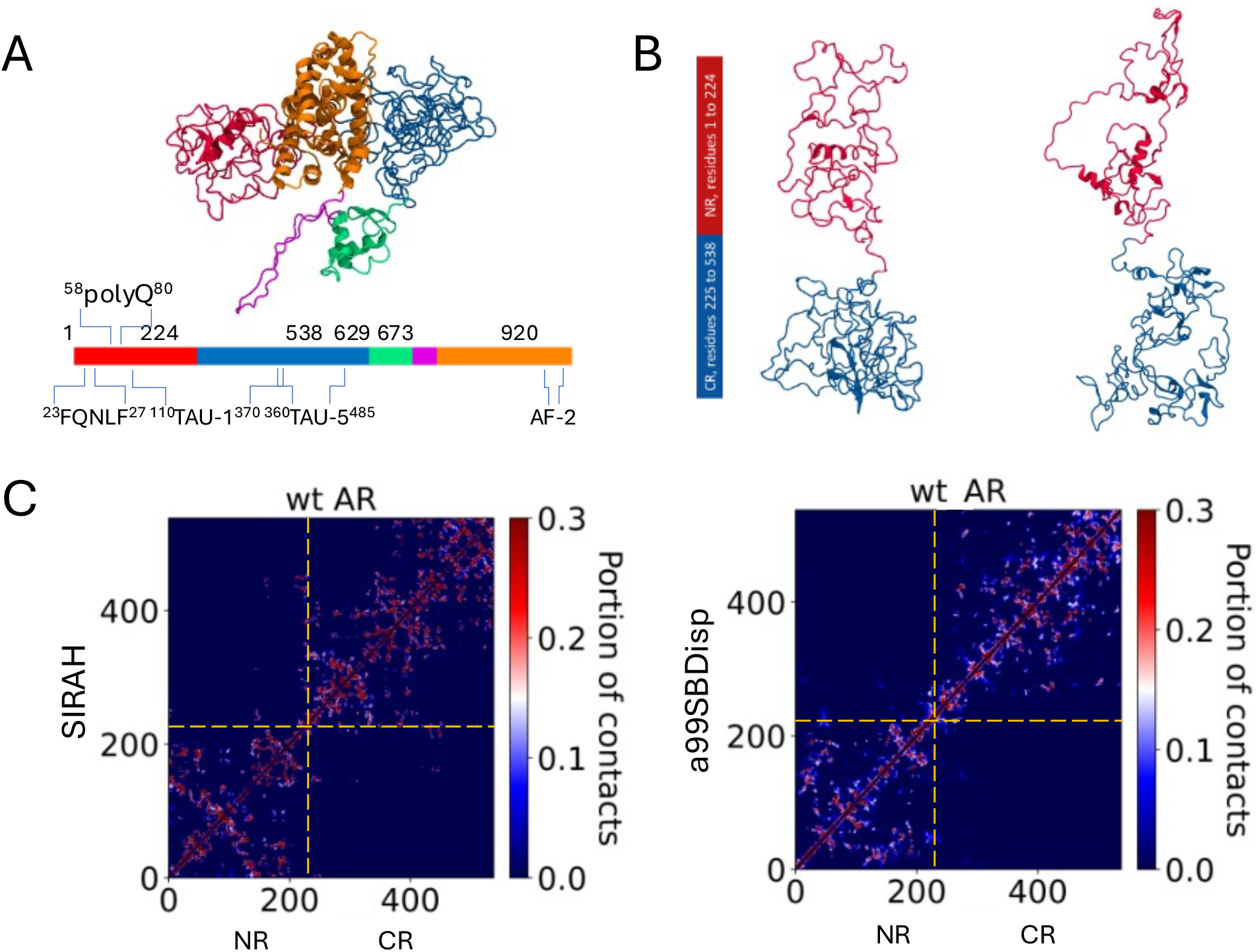
Wild-type AR-NTD forms into dynamic segregated subregions NR and CR (A) Full length model of AR-NTD color coded by region. red: NR; blue: CR; green: DBD; purple: hinge region and orange: LBD. Taken from Sheikhhassani et al (2022)^18^. (B) NR and CR subregion segregation in NTD is obtained with both coarse-grained SIRAH (left) and all-atom a99disp (right) forcefields. (C) Contact maps highlights dynamic regional segregation of NTD into NR and CR as seen both in SIRAH and a99SBDisp generated models.

pQ tracts occur frequently in the human proteome, predominantly in the intrinsically disordered regions of transcription factors^19^, yet their biological function is poorly understood^20^. One hypothesis is that the pQ tracts regulate activity by altering the stability of the complexes they form^21^ and that contractions and expansions have functional implications subject to natural selection^22^. Variations in length of pQ tracts have been linked to nine hereditary neurodegenerative diseases, including Huntington’s disease and SBMA, colloquially known as the pQ diseases^23^. The pQ tract in AR-NTD is also evidently involved in several other key AR pathologies. Apart from SBMA, variable lengths of this tract in AR have been linked to the onset and severity of prostate cancer^24,25^ and infertility^26^. The structural and functional consequences of pQ expansion towards the progression of SBMA is still a matter of debate with several hypotheses proposed. Some suggest that pQe AR-NTD are a neurotoxic species which lead to protein aggregate formation in the cytoplasm and nucleus^27^, while others have suggested that pQe AR-NTD itself is neurotoxic, affecting signaling functions^28^ and autophagy^29^, possibly leading to apoptosis. AR-NTD forms key protein-protein interactions (PPIs) with the other AR domains (DBD and LBD) as well with over 250 different proteins to exert its function^30^. Through intra-domain interactions with DBD, the NTD was previously shown to act as an allosteric regulator of DNA binding, while LBD interactions, also termed the N/C interaction, are seemingly important for transcriptional activity of AR. The interdomain interactions are with an extensive range of co-regulator proteins^31–35^. For example, RNA polymerase II-associating protein 74 (RAP74, a subunit of the TFIIF transcription factor) promotes transcriptional activities of AR while C-terminal of Heat Shock Protein (HSP)70 Interacting Protein (CHIP) mediates AR ubiquitination. While unknown, any mechanistic process through which pQe affects the interaction dynamics with these inter- and intradomain is plausible to contain important information regarding molecular mechanisms behind SBMA.

In this study, we employed extensive molecular dynamics (MD) simulations, topological analysis and advanced docking approaches to model the impact of pQe in AR-NTD (pQe: 45 glutamines, a length that has been associated with SBMA) compared to physiologically wild type (wt: 23 glutamines) present in healthy subjects. We investigated whether disease-related pQe tracts can alter conformational dynamics of NTD, monitored its consequences for interactions with DBD and co-regulators, and revealed how the pQe adds to aggregation propensity of AR in SBMA. We built a structural model that describes how the dynamic misfolding (a term we use to make a distinction with misfolding as it happens in structured proteins) of NTD leads to the gain/loss-of function of AR in SBMA.

## Results and Discussion

### Polyglutamine expansion alters local and global conformations and dynamics

We conducted all-atom MD simulations using a99SBDisp on isolated pQ tracts to understand the effect of expansion on their structural dynamics. Initially, we simulated pQ23 and pQ45 for 500 ns and found that pQ23 adopts an alpha-helical rod shape (Figure 2A), consistent with previous reports^36^. We observed from our simulations that pQ45 also adopts α-helical structures, albeit with different lengths and orientations. The intricate structural properties of pQ tracts, when isolated from their protein context, have been the subject of extensive scientific investigation and debate, and a variety of conformations have been observed, dependent on flanking amino acids^2^. These observations thus raise the question of how the expanded pQ tract impacts the global architecture and dynamicity of AR-NTD. To address this, we performed coarse-grained SIRAH MD simulations on full-length NTD with a pQe stretch, using our previously published AR modeling approach^18^. To ensure our initial conditions were independent of the results we performed these simulations multiple times (n=7) with different starting velocities. In contrast to wt-NTD (Figure 1), we observed that within 2 µs, the expanded pQ tract forms a globular structure different from what was observed for isolated pQ45 (Figure 2A), suggesting local misfolding upon expansion influenced by flanking residues. Importantly, the pQe-NTD forms a dynamic globular conformation whereby the segregation of the dynamic sub-regions is less prominently observed as in wt-NTD (Figure 2B). Furthermore, the pQ tract as well as the other physiologically important motifs, like the ^23^FQNLF^27^ and the KELCKAVSVSM (also known as the Androgen N-terminal Signature (ANTS) motif^37^: UniProt residues 235-245, pQe-NTD residues 257-267) remain surface exposed, despite the drastic architectural changes. We continued the simulations a further 3 µs (for all 7 runs) to follow its dynamicity over a longer time. The molecule showed high levels of structural dynamics, yet the dynamicity of pQe-NTD was reduced when compared to wt-NTD (Figure 2C), however, it still maintained a larger root-mean-square fluctuation (RMSF) when compared to a folded protein, like AR-LBD^38^.

**Figure 2.**
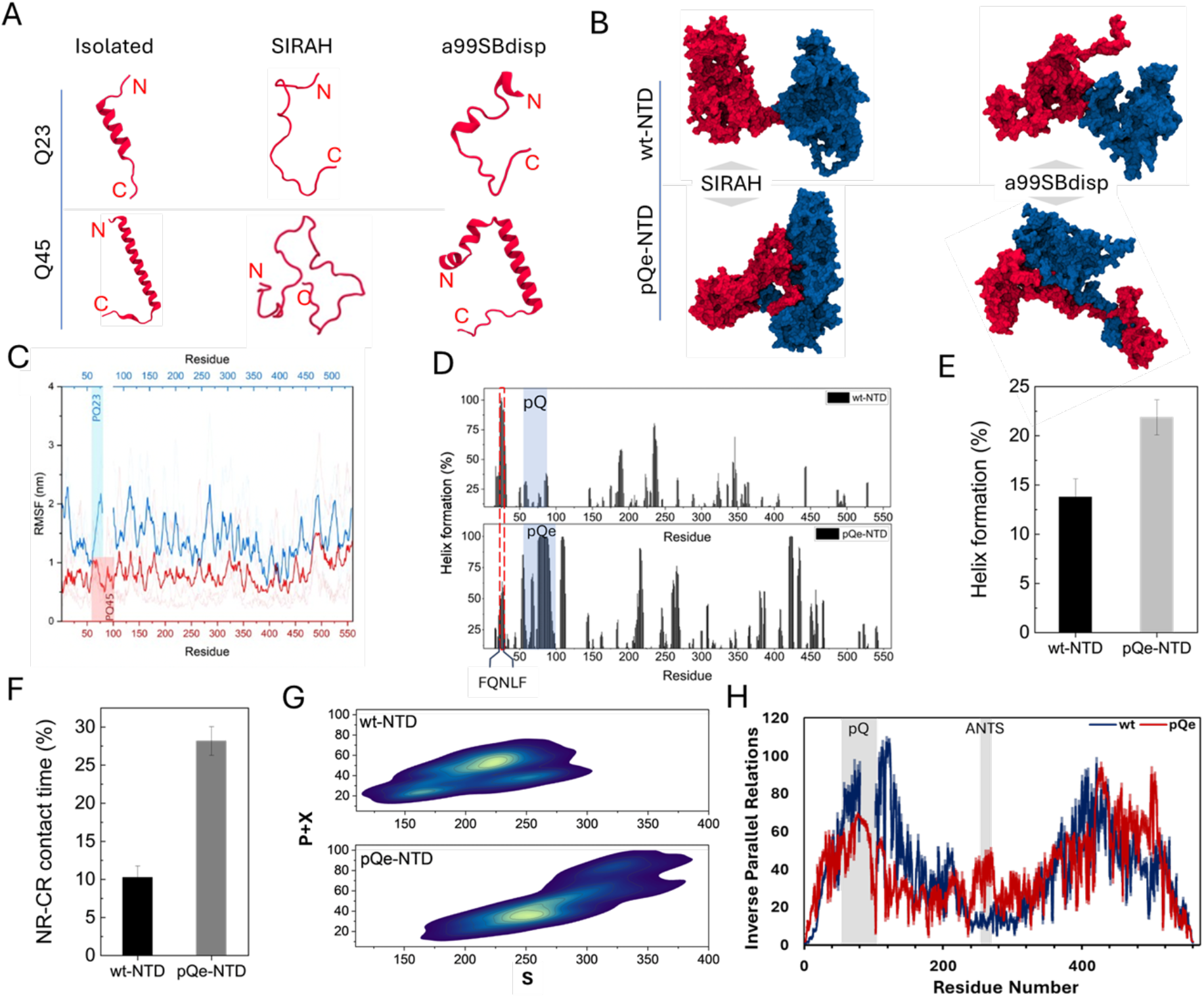
pQe results in local misfold and global alterations of protein dynamics (A) pQ tracts at the end of the simulations taken from “isolated simulations”, the SIRAH and a99SBDisp models. The results indicate that Q23 and Q45 tracts adopt different conformations. (B) pQ expansion results in an altered global conformation, both from SIRAH and a99SBDisp forcefields. (C) Root mean squared fluctuations (RMSF) calculated per residue indicates that pQ expansion reduces chain dynamics. (D) Residue-based α-helix analysis of wt- and pQe-NTD during the last 850 ns of all-atom MD simulations. Residues that are involved in a stable helical formation (probability >30%) are observed in the FQNLF region and sections of the pQ and pQe tracts. (E) Quantitative secondary structure analysis reveals that pQ expansion increases the α-helical content from 13.7 to 21.8% (error: STD over three simulations). (F) Contact time between the CR and NR regions of the wt-NTD and pQe-NTD. It is defined as the fraction of time during the last 850 ns of simulations when residues from the NR and CR were in contact. (error: STD over three simulations) (G) Two-dimensional density plots comparing the topological configuration space of the wt-NTD and pQe-NTD. Each contour plot shows the distribution of combined P+X topological relations as a function of S topological relations, aggregated over three independent simulation replicates. The contour levels represent the probability density of sampled configurations, with lighter shades indicating regions of higher probability. The axes represent the number of residues (× 1000) (H) Residue-based local circuit topology for WT and pQe AR NTD. The mean number of inverse parallel relations (± SEM, shown as shades) is shown as a function of residue number. The pQ tract and ANTS motif are highlighted by grey-shaded areas.

Given the importance of our observations, we decided to validate the identified conformational patterns of the NTD using highly accurate but computationally expensive all-atom simulations using a99SBdisp, a forcefield which is tailored to IDP modeling^39^. To avoid bias due to the choice of the starting conformations, we fully stretched our starting conformations and performed the all-atom simulations on stretched wt- and pQe-NTD. Our results on wt NTD are consistent with the SIRAH simulations, forming clearly segregated, highly dynamic, compact conformations on its N-terminal and C-terminal (Figure 1B) with no contacts between the two subregions (Figure 1C). Upon pQ expansion, the NR dynamically misfolds and interacts with the CR (Figure 2B and S1). This is reflected by an increased contact time between NR and CR (Figure 2F). In addition, as a result of the global restructuring caused by pQ expansion, the α-helical content in NTD increased by >8% as determined through structural analysis quantification (Figure 2D and E) and comparable to experimental observations^2^.

To understand the role of the glutamine residues in these altered fold dynamics, we mapped its interaction network with other NTD residues. It shows that the pQe tract formed longer-range contacts, including with residues in the CR region, a behavior not observed in wt-NTD (Fig S2). These long-range contacts also included directional hydrogen bonds, which likely explains the lack of regional segregation and reduced dynamicity in pQe. The CR residues that the pQ tract interacts with partially overlap with the ANTS motif (Figure S2)^40^.

To quantify the structural differences between wt and pQe AR NTD, we sought to capture key topological features of the protein chain. Unlike traditional geometric methods, topological analysis is particularly well-suited to quantify and categorize the residual structural properties of proteins that are continuously deforming, such as IDPs. By mapping the pQe NTD dynamics onto a topological space, we identified recurring motifs and patterns within the dynamics while showing a recognizable difference with wt NTD (Figure 2G). To probe these differences further, we tracked the topological dynamics of the NTD over time; employing the circuit topology (CT) framework^41–43^ to calculate the residue-based parallel (P), series (S), and cross (X) topologies within the chain’s conformations at 25 ns intervals (Figure S3). Residue contacts, defined by spatial proximity, serve as a fundamental topological representation of the chain, offering deep insights into the dynamics, and how they are altered in pQe. This revealed that, in agreement with our contact analysis, the pQ region in wt is enriched in inverse parallel relations (Figure 2G), indicating it is more sequestered in the structure than in pQe NTD. In contrast, the ANTS motif is more sequestered in pQe NTD (Figure 2G), revealing a long-range contact between the pQ tract and this motif in pQe NTD.

Overall, both SIRAH and a99SBDisp modelling support a model in which wt NTD adopt segregated dynamically compact regions. Importantly, local misfolding due to pQ expansion propagates to global reorganization in NTD, in which these segregated regions dynamically merge and unmerge. Next, we investigated the implications of the observed dynamic misfolding for inter-domain interactions.

### Gain of function, loss of regulation: pQ expansion induced dynamic misfolding alters protein-protein interactions

The roles of AR span genomic and non-genomic pathways^1^, which are both tightly regulated in cells. Our previous wt-AR-NTD model suggested a structural mechanism for the regulation of AR-DBD binding to DNA through competing NR and CR interactions. We showed that CR binding would cover the P-Box, the zinc finger of the DBD that binds to the major groove of DNA, and thus affect the ability of the DBD to interact with DNA^18^. These observations are consistent with previously published experimental studies, which identified the NTD and stretches of the CR as having DNA binding regulatory functions^44–48^.

To evaluate whether pQe affects interactions between the NTD and DBD, molecular docking analyses were conducted using both coarse-grained derived models, which offer broader conformational sampling, and all-atom derived models, which provide higher atomic accuracy but limited sampling. The coarse-grained results, which better capture the dynamic conformational flexibility of molecular interactions, have been prioritized in this study (Figure 3). In contrast, the all-atom results, offering finer structural resolution, are included in the supplementary information for additional validation (Figure S4).

**Figure 3.**
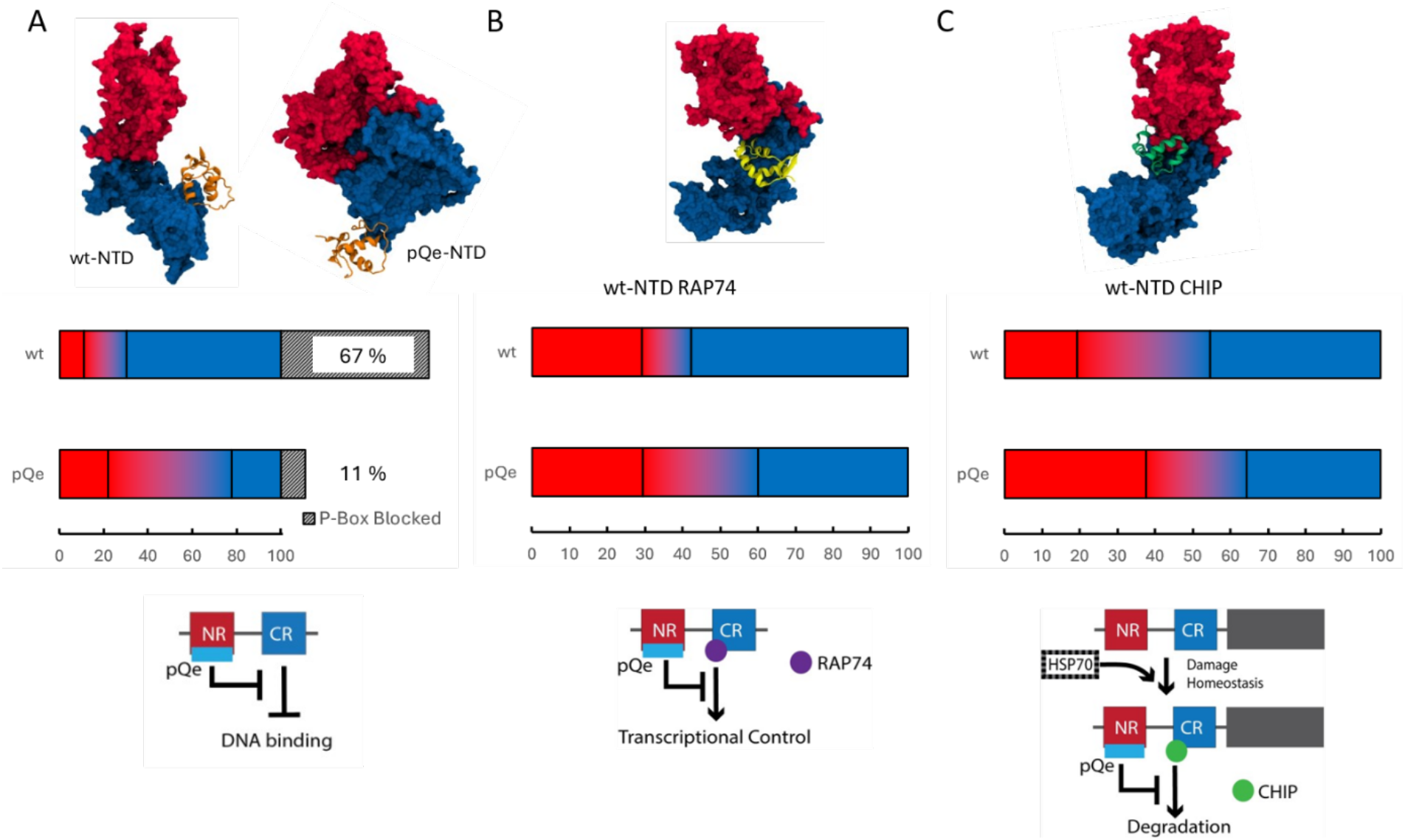
Protein-protein interactions are altered as a result of pQ expansion. (A) Representative structure of wt- and pQ-NTD interacting with DBD (orange). CR-mediated DNA binding regulation was reduced in pQe-NTD due to exposure of P-Box despite NTD-DBD interactions. (B) Representative structure of wt-NTD interacting with RAP74. pQ expansion induces a change in binding sites, reducing transcriptional control. (C) Representative structure of wt-NTD interacting with CHIP. Bar charts indicate the fraction of poses interacting with NR (red) NR/CR (dual colored) and CR (blue) residues. Graphics underneath highlight the mechanistic changes upon pQ expansion.

Our findings indicate that the altered structural dynamics of pQe-NTD significantly affect its interactions with the DBD. In the wild-type (WT) model, the majority of identified interacting residues localized within the CR region of the NTD. However, this interface was disrupted in the pQe variant (Figure 3A). Specifically, the ability of the NTD to block the P-box—a critical regulatory site for DNA binding—was markedly reduced, with only 11% of docking poses achieving P-box blockade compared to 67% in the WT model. This reduction aligns with prior reports of increased DNA binding in AR with expanded pQ tracts but contrasts with their observation of reduced AR transcriptional activity^49^.

To explore this apparent discrepancy, we used our docking approach to model interactions between pQe-NTD and RAP74, a subunit of the TFIIF complex^50,51^. RAP74 plays an essential role in the formation of the pre-initiation complex, facilitating the specificity and efficiency of RNA Polymerase II recruitment to DNA^52^. Consistent with the observed DBD interactions, our model revealed suppressed CR-mediated interactions with RAP74 in pQe-NTD (Figure 3B). These findings present a structural model illustrating how pQ expansion disrupts regulatory interactions critical to AR transcriptional activity.

### NTD has a propensity to aggregate as a result of altered dynamics by pQ expansion

Expansion of the glutamine tracts leads to ligand dependent protein aggregation in AR^53–57^, widely regarded as one of the pathological hallmarks of SMBA. To explore how the altered structural dynamics impact the dimerization of NTD, we first performed self-docking studies based on wt-NTD. Our dimer model demonstrates that the two monomers dynamically adopt both asymmetric head-to-tail (N-C) conformations (Figure 4A) and symmetric head-to-head (N-N) conformations (Figure S5A). These dimeric poses further reveal a spatial gap between the two molecules, in a manner that is consistent with the low-resolution cryo-EM AR structure^17^. This configuration supports the docking and interaction of the previously characterized active LBD dimer^58^ in agreement with the crystal structure of the AR-LBD dimer.

**Figure 4.**
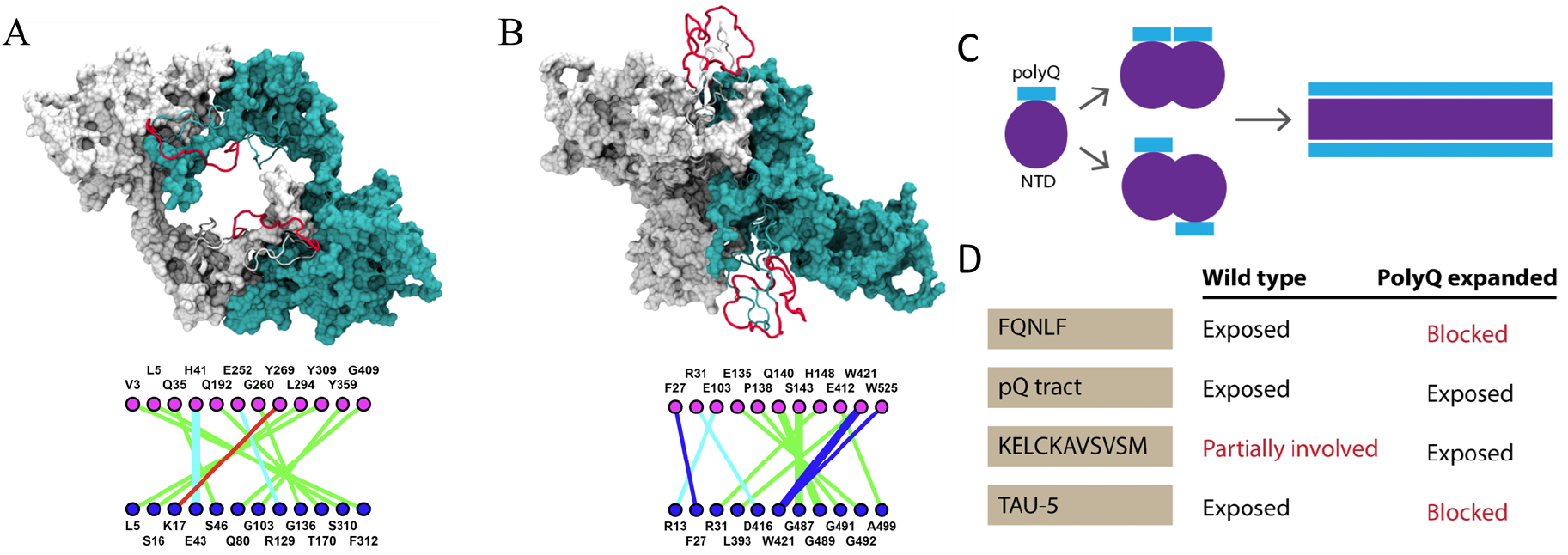
Multimerization of pQe NTD leads to fibrillar aggregates. Representative structure and residue interaction map of (A) wt-NTD and (B) pQe-NTD. The white and cyan structures represent one NTD monomer, the pQ tract is depicted in red. The precise interactions we saw between the two monomers for wt- and pQe-NTD are depicted under the representative structures. Different types of interactions are colored differently: Hydrogen bonds (green), Salt Bridge/Ionic (cyan), π-Cation (red), and π-Stacking (blue). (C) Aggregation model of pQe NTD, leading from a monomer to a fibril with the pQ tracts exposed. (D) Table of functionally important motifs and their involvement in a NTD dimer. “Exposed” means that the motif is exposed on the surface of the dimer, while “blocked” means that it is buried in the interaction interface. “Partially involved” means that a subpart of the motif is involved in the dimer interface.

Subsequently, we generated a pQe-NTD dimer, which illustrates a loss of the spatial gap (Figure 4B) conformational characteristics seen in the wild type dimer (Figure S5B). Notably, over 50% of the docking poses of pQe-NTD block the FQNLF motif (Figure 4B - interaction map and 4D), while the pQ tract and ANTS motif remain exposed on the surface of the dimer. This last observation shows strong similarities with studies on pQe Ataxin-3, a protein associated with Machado–Joseph disease, another polyglutamine expansion disorder. In Ataxin-3, the expansion of the pQ tract leads to misfolding and aggregation in a two-step process, transitioning from amyloid-like fibers to insoluble aggregates^59–62^.

Oligomerization beyond dimers has been suggested for both healthy and pathogenic isoforms of AR^58,63^. Previous reports have shown that pQe-AR-NTD undergoes an altered oligomerization process compared to wt-NTD, forming fibrils instead of annular oligomers^64^. To explore if our pQe-NTD model could provide atomistic detail to this process, we built a higher-level aggregate model, factoring in tetramer and octamer pQe NTD models (Figure S6). Our results mirror those observed in dimerization, where the pQe tracts are exposed on the surface, while the functional areas like the Transactivation Unit (Tau)-5 and FQNLF, mediate interactions between individual molecules. Oligomerization to an octamer demonstrates a clear tendency to form a fiber, in agreement with the AFM results^64^. This data allowed us to build a model of multimerization of pQe-NTD (Figure 4C), from a monomer into oligomers.

The behavior of pQe-AR observed here aligns with previous reports on polyglutamine expanded proteins and provides a foundational basis for a unifying mechanism for polyglutamine and other misfolding diseases. In these disorders, (toxic) monomeric proteins aggregate into insoluble fibers, a process initially thought to underlie pathogenicity^65^. However, other studies have questioned whether aggregation is the primary cause of cytotoxicity^66^.

Could the formation of insoluble fibers be cytoprotective? Previous reports on Huntingtin (Htt) suggest that inclusion body formation reduces the diffusion of Htt protein, thereby decreasing neuronal cell death^67^. Our model, where the FQNLF and other functional motifs are blocked through multimerization may suggest a similar protective mechanism. The FQNLF is crucial for AR activation, a process partly regulated by the competition between chaperone proteins such as HSP40 and HSP70, and the AF-2 coactivator binding pocket. NMR data has shown that HSP70 directly interacts with FQNLF in wt-NTD, preventing aggregation *in vitro*^68^. Furthermore, modulating HSP70 was found to increase solubility, ubiquitination, and clearance of pQe-NTD. Hsp70 cycles between ATP- and ADP-bound states, with the ADP-bound state exhibiting significantly enhanced substrate binding affinity^69^. This state allows HSP70 to discriminate between properly folded and damaged proteins^70^. The most effective HSP70 compounds stabilized it in the ADP-bound states, which suggests that, in the case of pQe-AR, HSP70 wrongly releases AR-NTD.

To explore chaperone interactions further, we examined the binding of HSP70 and CHIP with both wt NTD and pQe NTD. We observed that while HSP70 interacts through the region with the FQNLF in wt-NTD, as reported in the NMR data, this interaction is absent in pQe-NTD (Figure S7). While CHIP interacted with wt-NTD primarily through CR residues, this interaction was significantly reduced in pQe-NTD (Figure 3C). Together these results suggest that in the absence of HSP70 modulation, pQe-AR is less likely to be soluble and ubiquitinated for degradation (Figure 3C, model). Multimerization of pQe AR leads to aggregation into insoluble fibers, which, at least initially, may as a cytoprotective mechanism.

## Conclusions

This study presents the first molecular model of expanded polyglutamine tract containing NTD of AR, offering unprecedented insights into the structural dynamics and functional implications of pQ expansion. Using extensive *in silico* simulations, topological mapping, and molecular docking analyses, we have provided detailed understanding of the complex intra- and interdomain interaction within AR-NTD, highlighting the effects of pQe on its structural reorganization, interactions with other AR domain and cofactor binding, particularly RAP74, HSP70 and CHIP.

Our findings reveal that expansion of the pQ tract induces a global dynamic restructuring of the AR-NTD, driven by long-range interactions of the glutamine residues. This reorganization disrupts the native dynamic regional segregation seen in wt AR-NTD. Importantly, these long-range interactions seem to include the ANTS motif, which was previously reported to crosstalk with the pQ tract of AR in the control of aggregation^40^. Interestingly, that study demonstrated that fibrillar aggregation of NTD fragments only occurred with pQe tracts, suggesting that the distance between the glutamine tract and ANTS was of importance. Our pQe NTD model does provide topological information on molecular interactions between the pQe and the ANTS motif. In contrast, this interaction is not seen with the wt-NTD. This solidifies the potential importance of this motif and the positive regulatory crosstalk these have.

Additionally, our model underscores the delicate balance of protein-protein interactions (PPIs). While pQ expansion results in the disruption of critical interactions – such as those between AR and TFIIF through RAP74 – leading to impaired gene regulation, it may simultaneously enhance DNA binding through the loss of self-inhibitory interactions. Furthermore, our results suggest that aggregation of pQe-AR could represent a cytoprotective mechanism. However, this aggregation disrupts key PPIs, including interaction with HSP70 and CHIP, which are essential for protein homeostasis. These alterations in PPIs as a result of pQe mediated dynamic misfolding can ultimately disrupt numerous cellular processes essential for cell survival, from homeostasis to gene expression. Dysregulation of these multiple pathways could culminate in toxicity in a cumulative fashion, suggesting that therapeutic targeting of a single pathway may not provide complete amelioration. In contrast, reports of reducing cellular levels of pQe AR have emerged as an appealing therapeutic strategy to target the proximal mediator of disease pathogenesis^68^. In this regard our model opens new therapeutic avenues by revealing the distinct interaction patterns between the pQe tract and AR motifs. Targeting these interactions could potentially stabilize or reverse the pathological effect of pQe-AR. Of course, further experiments will be necessary to gain a deeper understanding of the altered transcription and regulation through PPIs by AR-NTD. Preliminary findings (Supplementary material) suggest that the interaction between the pQ tract and the LBD may further influence AR function by modulating the N/C interaction^71^, which can be affected by neighbouring motifs^72^, Although current computational methodologies limit the study of interactions between disordered regions like NTD and folded domains like LBD, our initial peptide docking results provide intriguing evidence that the pQ tract interacting with a pocket on the LBD (Figure S8) could modulate the FQNLF/AF-2 interaction through a conformational change (Figure S9). This binding pocket was previously identified as a highly conserved structural pocket, the BF-3^73^. It is possible that expansion of the pQ tract affects this process in SBMA.

In conclusion, despite the incredible complexity of modelling nuclear receptors and large intrinsically disorder proteins, our approach has generated atomistic-level insights that are consistent with experimental observations. This work not only provides a detailed understanding of the structural dynamics of AR in the context of polyglutamine expansion but also sets the stage for future studies of other disordered proteins and their pathological mutations. By unlocking new mechanistic insights, this model paves the way for the development of novel therapeutic strategies aimed at mitigating the effects of misfolding and aggregation in neurodegenerative diseases.

## Methods

### Molecular dynamics simulations of AR-NTD

This study presented MD simulation data on wt and pQe-NTD using SIRAH and a99SDisp. All details of our simulations are depicted in Figure S10. Initial representations of pQe AR-NTD were generated using I-TASSER^74^ and mapped to the coarse-grained (CG) SIRAH force field. The CG models were solved in octahedron boxes with a minimum distance of 1.2 nm from the solute using pre-equilibrated WT4 water molecules. The ionic strength was set to 0.15 M by randomly replacing WT4 molecules with Na^+^ and Cl^−^ CG ions. The system was prepared following standard SIRAH protocols. First, energy minimization was performed with positional restraints of 1000 kJ mol^−1^ nm^−2^, followed by a 100 ps NVT simulation at 300 K and a 100 ps NPT equilibration. Production simulations were performed in the NPT ensemble at 310 K and 1 bar using GROMACS 2020^75^ with a time step of 20 fs. PME electrostatics were used with the Verlet scheme and a cut-off of 1.2 nm. Solvent and solute were coupled separately to velocity rescale thermostats with coupling times of 2 ps. Pressure was controlled by the Parrinello–Rahman barostat with a coupling time of 10 ps. Production trajectories were generated for 5 µs.

### Initial structure preparation for all-atom simulations

Initial structures for all-atom simulations were generated from the I-TASSER structure. Initially, structures were subjected to pulling via steered MD simulations to form a fully extended chain. Subsequently, 50 ns all-atom simulation in implicit solvent were conducted to produce new initial structures. The QuickMD module of Visual Molecular Dynamics (VMD)^76^ in conjunction with the NAMD molecular dynamics package^77^.

### All-atom molecular dynamics simulations

All-atom simulations on wt-NTD, pQe-NTD and isolated pQ tracts (Q23 and Q45) were performed using a99SDisp forcefield with compatible amber water molecules. A total of 1000 ns were simulated with a 2 fs time step. Neighbour searching was performed every 10 steps. The PME algorithm was used for electrostatic interactions with a 1 nm cut-off. A single cut-off of 1.006 nm was used for Van der Waals interactions. Temperature coupling was done with the V-rescale algorithm and Pressure coupling was done with the Parrinello-Rahman algorithm. Trajectory analysis and visualization for all simulations were performed using VMD and GROMACS tools.

### Circuit Topology analysis

Circuit topology parameters were analyzed from the final 850 ns of each aa99SBdisp simulation using custom Python scripts. These scripts were modified to include energy and length filtering, as well as circuit decomposition. Residue contacts were defined by a 4.5 Å distance cutoff, requiring at least five atoms from each residue to be within this distance. To avoid local overcounting, the three nearest neighbors of each residue were excluded from the contact analysis.

Residue-based local circuit topology was extracted using MATLAB based on established methods^42,43,78^. A cutoff scheme of 5 or more atom-atom contacts within 4.5 Angstroms was employed, excluding contacts within 3 residues along the protein sequence. Circuit topology was calculated over the final 700 ns of the all-atom simulations, at a 25 ns timestep, for all three simulations. Error was calculated as SEM over all extracted frames.

### AR-NTD protein-protein interactions

The atomic coordinates of the pQe-NTD and wt-NTD were taken from our structural SIRAH and aa99SBdisp models and used as input for protein-protein interactions using biased rigid body docking with ClusPro 2.0 webserver^79^. The atomic coordinates of partner proteins (AR-DBD: PDB 1R4I; AR-LBD: PDB 1T7T; CHIP: residue 311-369 extracted from AlphaFold ID P50502; RAP74 C-terminal domain: PDB 1I27; HSP70: 4PO2) were selected as ligands. For multimeric docking, the atomic coordinates of our NTD models were taken as both receptor and ligand.

All poses predicted by ClusPro were inspected using VMD and analysed using a custom-made Python script; utilizing different libraries: madplotlib^80^, mdanalysis^81–83^, mdtraj^84^, numpy^85^, pandas^86^, scipy^87^ and seaborn^88^.

## Author contributions

L.W.H.J. Heling: Investigation, Data Curation, Visualization, Writing – Original Draft, Writing – Review & Editing

V. Sheikhhassani: Investigation, Methodology, Data Curation, Visualization, Writing – Original Draft, Writing – Review & Editing

J. Ng: Investigation, Funding Acquisition, Writing – Review & Editing

M. van Vliet: Investigation

A. Jiménez-Panizo: Investigation, Writing – Review & Editing

A. Alegre-Martí: Investigation, Writing – Review & Editing

J. Woodard: Investigation, Writing – Review & Editing

W. van Roon-Mom: Writing – Review & Editing

I. J McEwan: Investigation, Resources, Writing – Review & Editing

E. Estébanez-Perpiñá: Investigation, Resources, Writing – Review & Editing

A. Mashaghi: Conceptualization, Investigation, Project Administration, Supervision, Writing – Original Draft, Writing – Review & Editing

## Data availability

The raw simulation data generated during this study have been deposited in the following Zenodo repository: DOI 10.5281/zenodo.14512908

## Acknowledgements

The authors thank Dr. Annemieke Aartsma-Rus for discussions.

