## Supplementary Information for "Polyglutamine expansion induced dynamic misfolding of Androgen Receptor"

**Supplementary materials: Wild-type pQ/LBD interaction analysis**

A. pQ20 interacts with BF3 pocket in LBD which may promote FQNLF/AF-2 interactions

In order to understand the impact of pQ expansion on the interaction between NTD and LBD of AR (termed the N/C interaction), we first explored the behaviour of wt-PQ in this process. Thus, we investigated the interaction between isolated pQ20 and AR-LBD (PDB: 1T7T) using docking analysis. Our analysis revealed three distinct interacting regions on the surface of the protein, located mainly between amino acids 670-730, 770-795, and 830-900 (Fig S4A). Of these, the first region, previously identified as the BF-3 region, interacts with the main cluster of pQ20 poses (Fig S4B), composed of 12 members, and was thus selected for additional interaction analysis. Our findings showed that seven distinct amino acids on the LBD surface are involved in the peptide-protein binding interface, involving both polar and nonpolar contacts with an almost equal interface area (Fig S1B). Furthermore, two hydrogen bonds were observed between the side chain of Asn833 and the main chain of Gly724 with two glutamine residues (Gln 1 and 4) in the peptide structure, which may contribute to stabilizing the pQ20-LBD interactions.

To investigate possible conformational changes induced by pQ20 binding to BF-3, we performed an all-atom MD simulation for 300 ns using the best pQ20 pose in complex with LBD as the initial structure. Interestingly, LBD showed almost the same RMSD changes over the simulation time, independent of pQ20 interaction, indicating that global conformational dynamics of LBD did not alter upon interaction with pQ20. However, higher average RMSF values were recorded in three regions, amino acids 690-695, 770-774, and 855-857, upon pQ20-LBD interaction. These residues are in helices 3, 5 and 9, respectively. Finally, representative structures of AR-LBD and pQ20-LBD complex were superimposed to examine the structural differences (Fig S5). Several structural changes were identified in LBD due to the pQ20 binding, compared with LBD in isolation. In particular, helices number 4, 5, and 9 rotated 3° and helix 12 6° with respect to their original positions. Moreover, the area of the AF-2 groove was expanded by moving helices 3, 4, 9, and 12 away from the groove, accompanied by moving the helix 5 towards the groove, which kept the general configuration of the AF-2 intact. Overall, our findings provide insights into the structural basis of pQ20-LBD interactions and potential conformational changes induced by this interaction.

B. Methodology: AR-LBD interactions analysis with polyQ tract

AR-LBD structure in complex with its natural ligand was available through the protein data bank. Prior to performing molecular docking, a total of 100 ns were simulated with a time step of 2 fs for relaxing the AR-LBD structure. All MD simulations were carried out using GROMACS package. Neighbour searching was performed every 20 steps. The PME algorithm was used for electrostatic interactions with a cut-off of 1.2 nm. A single cut-off of 1.221 nm was used for Van der Waals interactions. Temperature and pressure coupling were done with the V-rescale and Parrinello-Rahman algorithms respectively.

Due to the technical limitation of peptide-protein interaction analysis algorithms, which accept a maximum of 20 residues for peptide structures, we performed wt pQ-LBD interactions using a truncated length of 20 Q residues. In this study, we used pepATTRACT[1], a flexible peptide-protein docking approach in ATTRACT, to find the interaction sites of the wt-polyQ tract (Q20) with AR-LBD. We used the sequence of the peptide as the initial input for the peptide structure. Despite these constraints, to broaden our analysis, we overcame the limitations by employing the ClusPro rigid body protein-protein docking algorithms to dock Q23 against LBD. Remarkably, the highly ranked poses from ClusPro align well with the results obtained from pepATTRACT. However, acknowledging the higher precision of flexible docking algorithms in predicting residue coordinates at the binding interface, we specifically selected the top 50 highest-ranked peptide poses derived from pepATTRACT for further detailed analysis. The energy-minimized structure of the LBD in both analysis was used as the receptor protein.

### Supplementary Figures

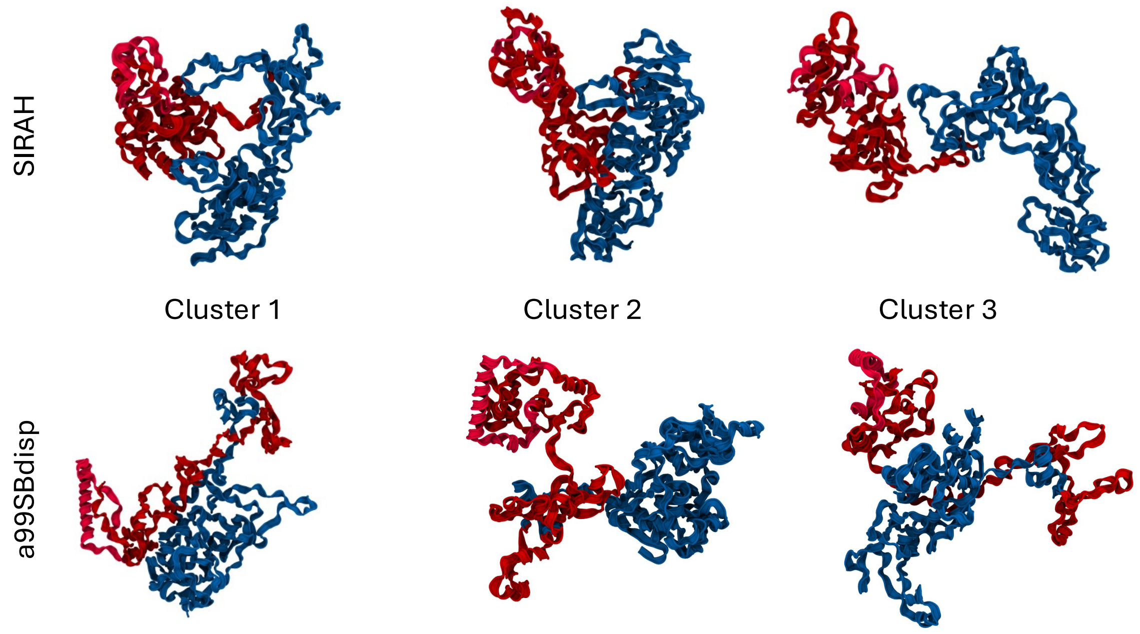


*Figure S1 Three largest clusters from SIRAH and all-atom a99SBdisp simulations. NR and CR residues are depicted in red and blue respectively.*

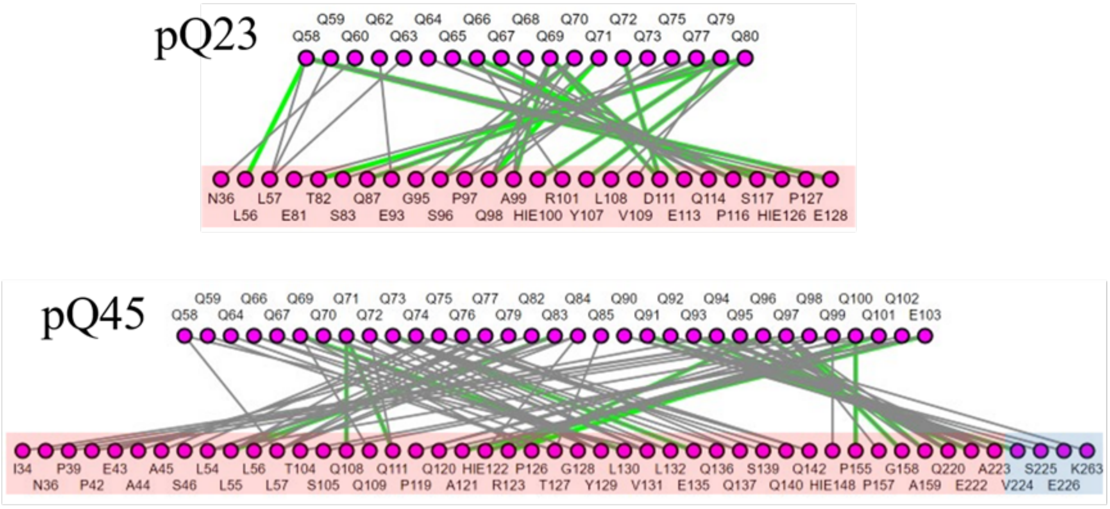


*Figure S2 Interaction network of the polyQ tracts in wt-NTD and pQe-NTD with other NTD residues. NR residues and CR residues are depicted with red and blue boxes respectively. In this interaction analysis hydrogen bonds are shown in green and close proximity contacts are colored in grey.*

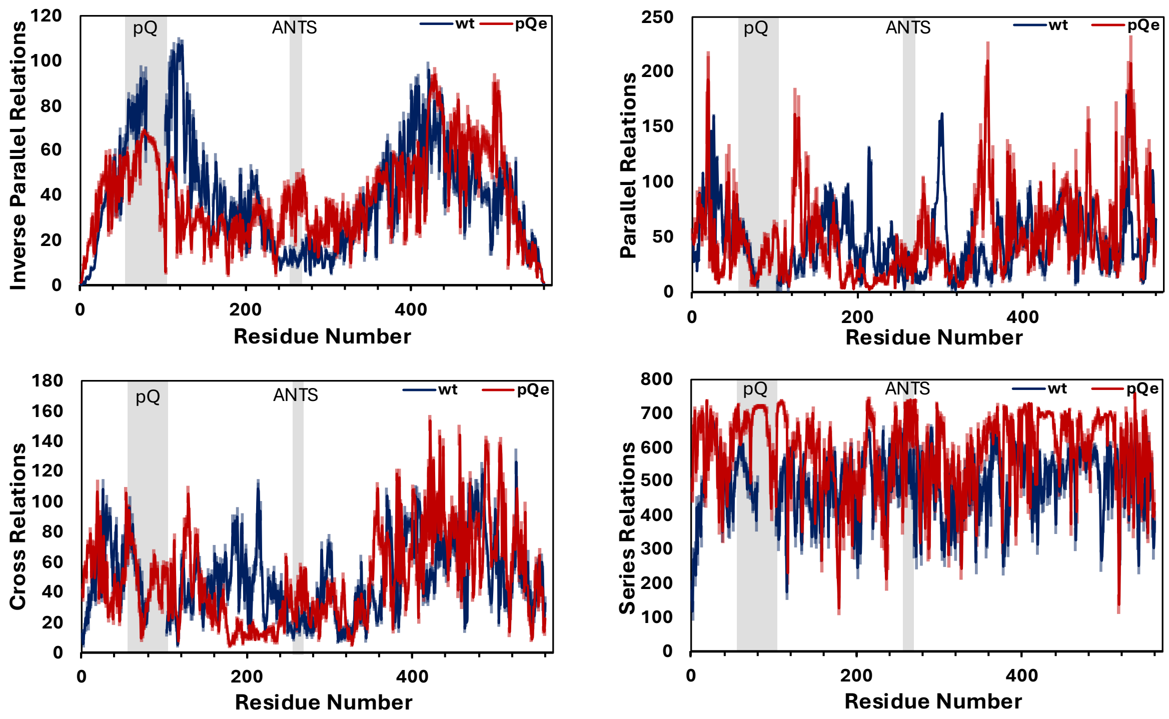


*Figure S3 Residue-based local circuit topology for wt and pQe AR-NTD. Shown are mean number (± SEM, depicted as shades) of inverse parallel (IP), parallel (P), cross (X), and series (S) relations as a function of residue number. The pQ region and ANTS motif are highlighted by the grey boxes.*

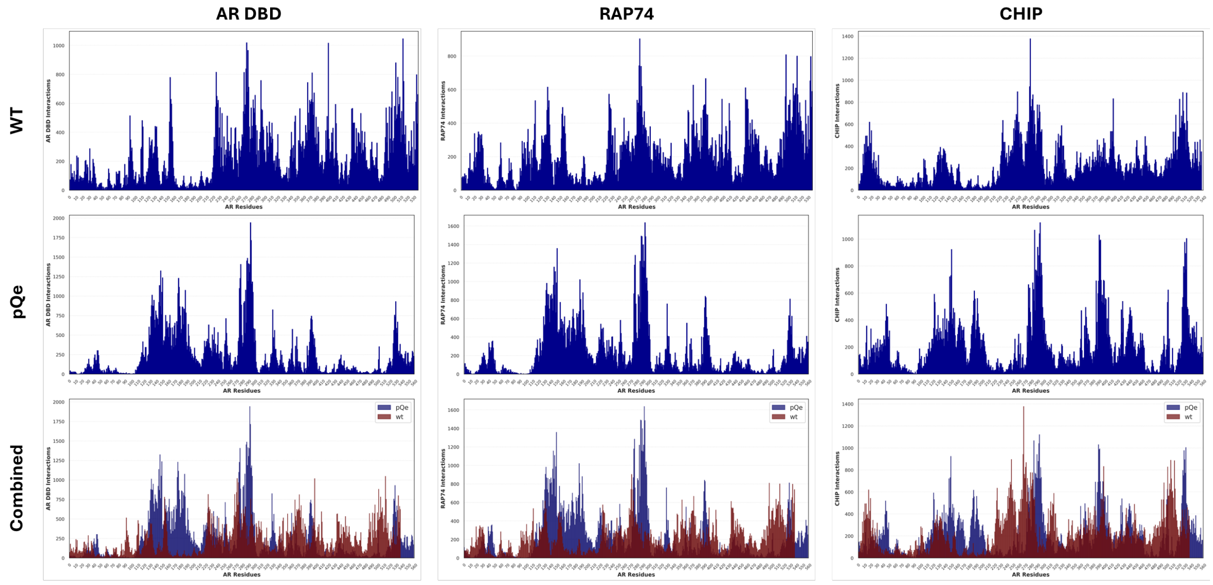


**AR DBD**

| | | NR | NR/CR | CR | P-box Blocked |
|---|---|---|---|---|---|
| SIRAH | wt | 11.1 % | 69.5 % | 19.4 % | 66.7 % |
| | pQe | 22.2 % | 22.2 % | 55.6 % | 11.1 % |
| a99SBdisp | wt | 22.8 % | 38.6 % | 38.6 % | 59 % |
| | pQe | 0.9 % | 88.9 % | 10.2 % | 0.9 % |

**RAP74**

| | | NR | NR/CR | CR |
|---|---|---|---|---|
| SIRAH | wt | 29.1 % | 13.3 % | 57.6 % |
| | pQe | 29.4 % | 30.8 % | 39.8 % |
| a99SBdisp | wt | 21 % | 27.2 % | 51.8 % |
| | pQe | 1.8 % | 87.1 % | 11.1 % |

**CHIP**

| | | NR | NR/CR | CR |
|---|---|---|---|---|
| SIRAH | wt | 19.5 % | 35.1 % | 45.4 % |
| | pQe | 37.5 % | 26.7 % | 35.8 % |
| a99SBdisp | wt | 16.1 % | 43.5 % | 40.4 % |
| | pQe | 5.9 % | 77 % | 17.1 % |

*Figure S4 wt and pQe interaction histograms with DBD, RAP74 and CHIP based on ClusPro using our aa99SBdisp derived models. Height of the bars indicate the total interaction count of that residue. The tables below show the percentages of exclusive interactions with NR and CR or both.*

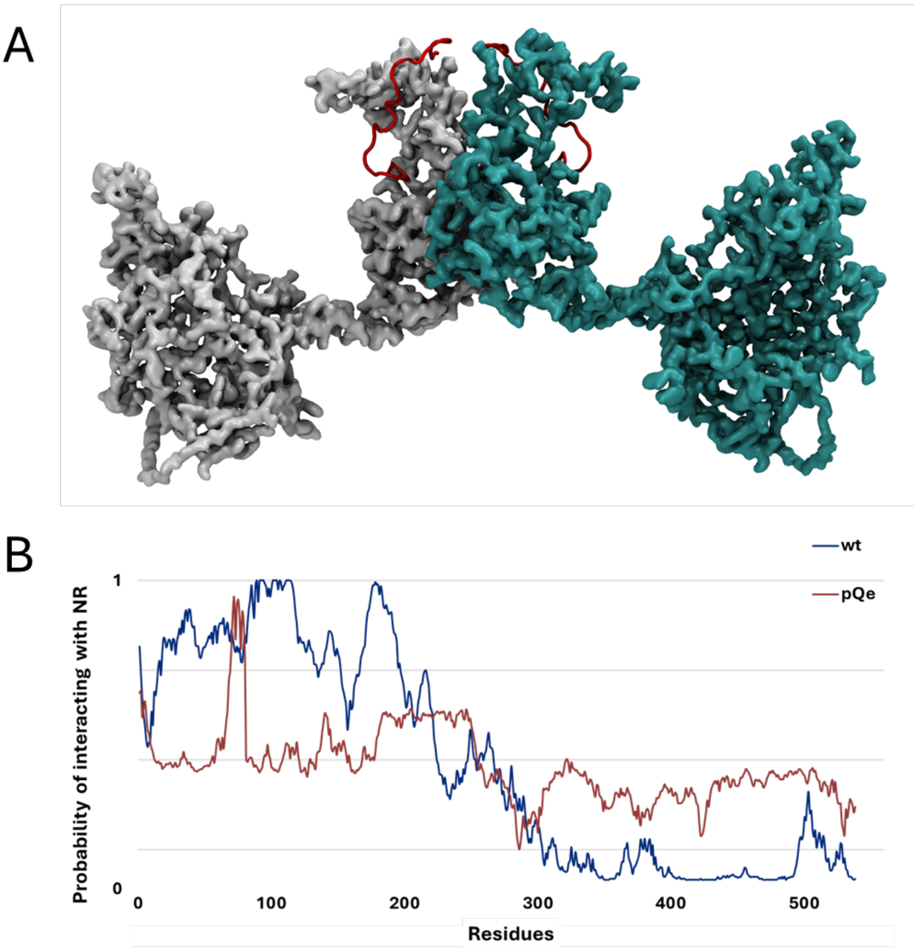


*Figure S5. AR dimerization is altered in pQe (A) wt-wt dimer (based on CG models) demonstrating a head-head conformation. (B) Residue based probability of interacting with NR residues in the other AR molecule in the dimer (based on all-atom derived models). In wt, the NR shows a high probability of interacting with the NR of the other monomer, and a lower probability to interact with the CR. This trend is lost in pQe, showing a relatively similar probability across the chain.*

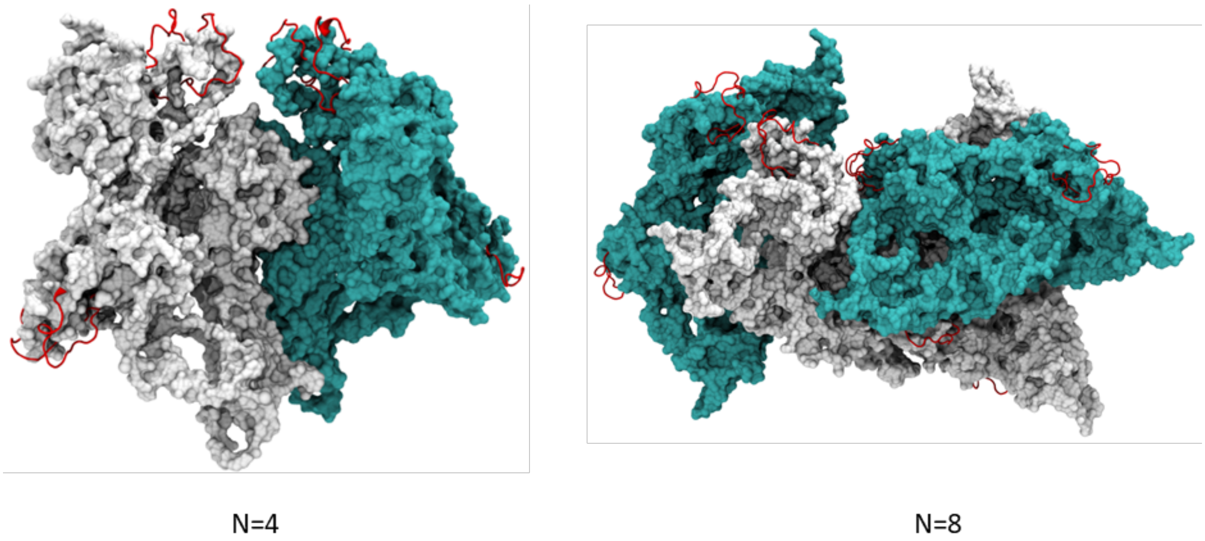


*Figure S6. Oligomerization of pQe NTD continues the trend of covering functional motifs and exposing pQ tracts. (A)Tetramer and (B) Octamer of pQe-NTD where white and cyan each represent two NTD molecules. The pQ tracts are depicted in red. N= number of NTD monomers in the structure.*

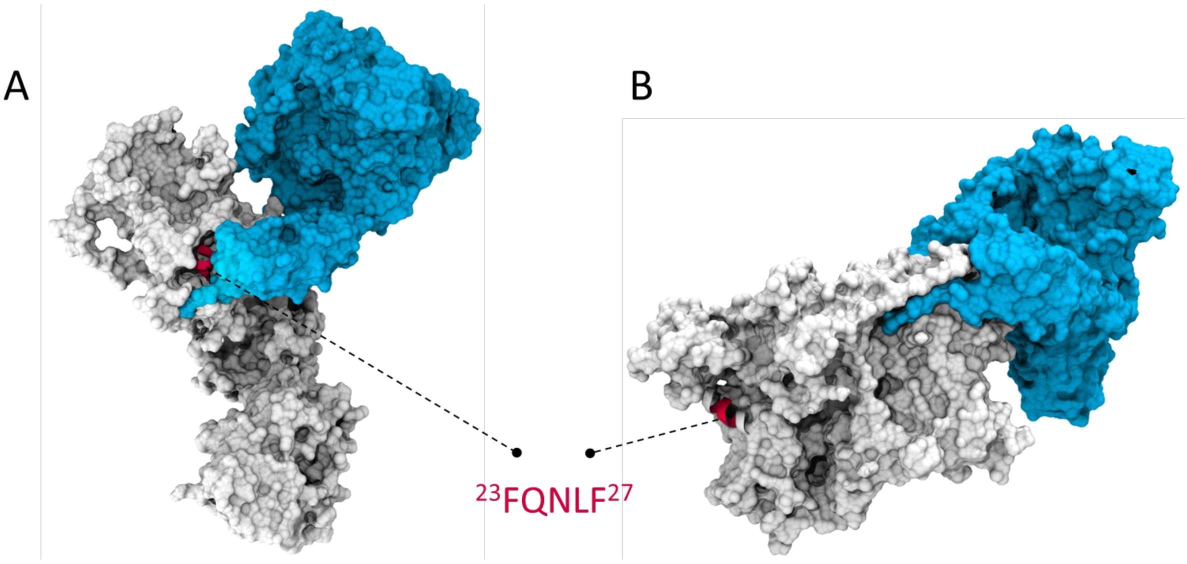


*Figure S7. HSP70 interactions with NTD are altered upon pQ expansion. Representative structures of (A) wt-NTD and (B) pQe-NTD in white docked using ClusPro with HSP70 (Blue; structure from Alphafold: AF-P11142). The interaction with the FQNLF motif (depicted in red) seen in wt-NTD is lost upon pQ expansion. It is possible that the loss of this interaction is the basis for reduced solubility of AR in SBMA.*

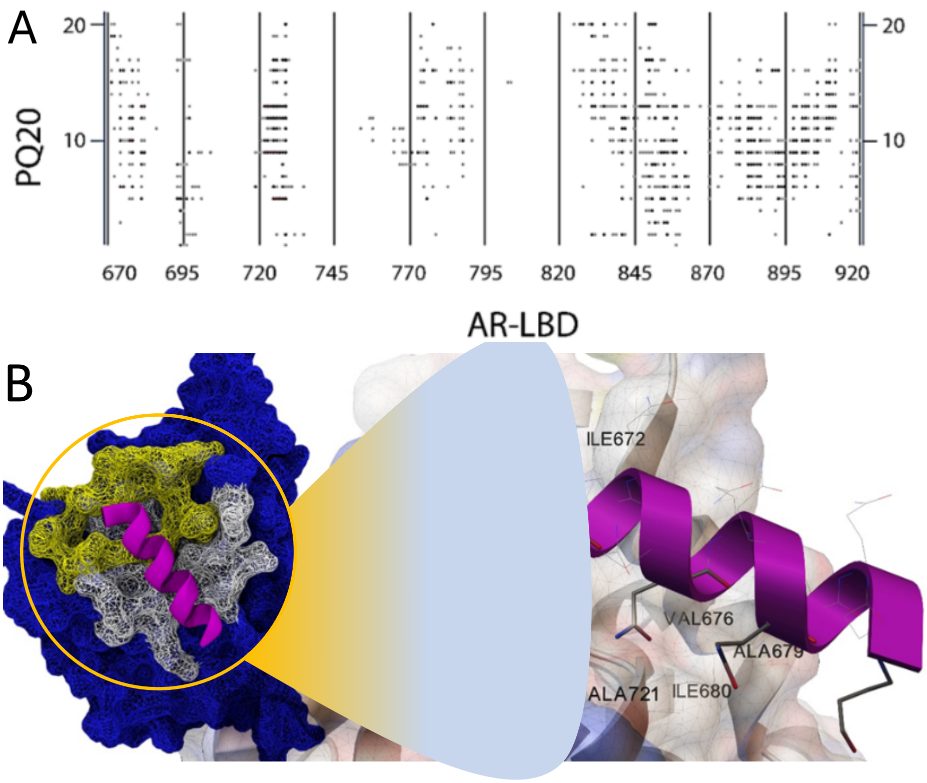


*Figure S8. The pQ tract binds to the LBD through the BF-3 pocket. (A) Contact map analysis of interactions between PQ20 and AR-LBD. (B) interacting region of PQ20-AR-LBD (white) shows a considerable surface overlap with BF-3 pocket (yellow). Interacting residues on the surface of AR-LBD. Small spheres in green shows hydrogen bonds formed between GLN residues of peptide and ASN 833 and GLY 727.*

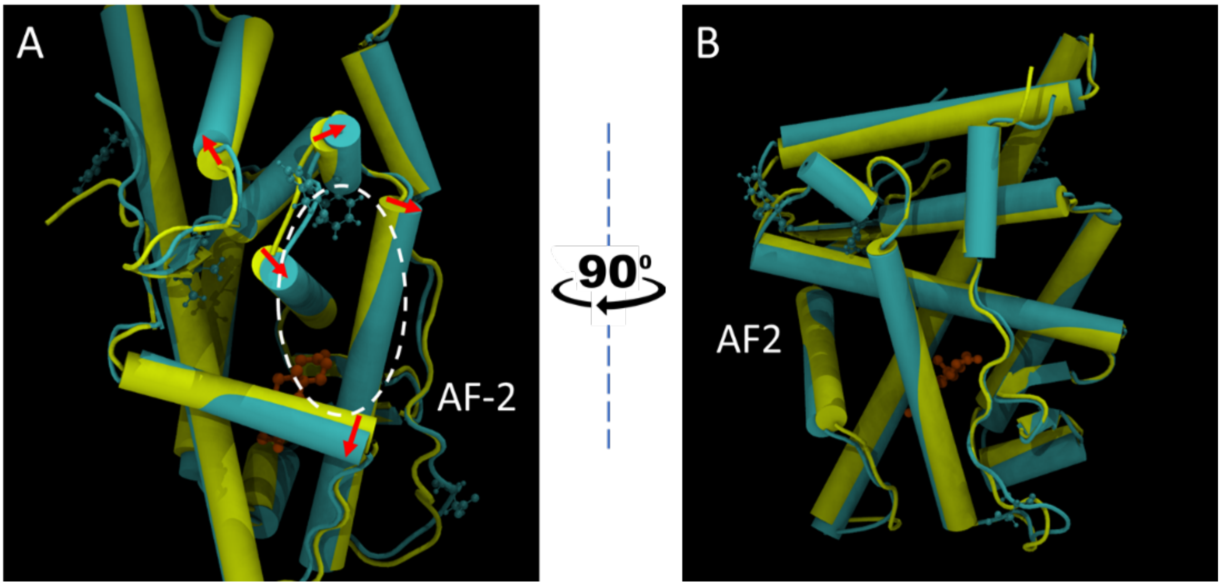

*Figure S9. pQ interactions with BF-3 induce conformational change in LBD, opening the AF-2 pocket. (A) Superimposition of representatives of PQ20-LBD and AR-LBD in isolation from AF-2 view and (B) from side view by 90 clockwise rotation. AR-LBD is drawn in yellow, and LBD structure isolated from PQ20-LBD complex in cyan. The red arrows on (A) mark the local structural changes between the two structures*

| | Force Field | Time | Starting Condition | Runs |
|---|---|---|---|---|
| wt-NTD | SIRAH | 5 µs | I-TASSER | 3 |
| wt-NTD | a99SBDisp | 1 µs | Stretched chain | 3 |
| pQe-NTD | SIRAH | 5 µs | I-TASSER | 7 |
| pQe-NTD | a99SBDisp | 1 µs | Stretched chain | 3 |

*Figure S10. Table of the structures and subsequent simulations with forcefields, times and starting conditions.*
